# Distinct forms of structural plasticity of adult-born interneuron spines induced by different odor learning paradigms

**DOI:** 10.1101/2023.07.20.549882

**Authors:** Aymeric Ferreira, Vlad-Stefan Constantinescu, Sarah Malvaut, Armen Saghatelyan, Simon V. Hardy

## Abstract

During development and in adulthood the morpho-functional properties of neural networks constantly adapt in response to environmental stimuli and learned experiences. One of the processes that allows neuronal networks to be constantly reshaped is synaptic plasticity, which is induced in response to sensory experience and learning. Synaptic plasticity allows for the formation/elimination of synaptic connections as well as the strengthening of pre-existing ones. The olfactory system is particularly prone to constant morpho-functional reshaping of neural networks and synaptic rewiring throughout the lifespan of an animal, mainly because of the presence of continuous neurogenesis in the olfactory bulb (OB). This constant synaptic rewiring brought by adult-born neurons is modulated by the level of odor-induced activity and olfactory learning. It remains, however, unclear whether the complexity of distinct odor-induced learning paradigms and sensory stimulation induces different forms of structural plasticity. In the present study, we developed an analytical pipeline to perform 3D reconstructions of spines from confocal images followed by clustering of reconstructed spines based on different morphometric features and in relationship with different sensory stimuli and learning paradigms. We show that while sensory deprivation decreased the overall density of adult-born neurons in the OB without any noticeable changes in the morphometric properties of these spines, simple and complex odor learning paradigms triggered distinct forms of structural plasticity. A simple odor learning task affected the morphometric properties of the spines without any changes in spine density, whereas a complex odor learning task induced changes in spine density, without substantial changes in the morphology of the spines. Our work reveals the vast panoply of distinct forms of synaptic plasticity of adult-born neurons in the OB tailored to the complexity of odor-learning paradigms and sensory inputs.

## Introduction

During the early postnatal period, the developing brain is shaped and remodeled in response to environmental stimuli. The major changes that occur during these critical periods of development, which are marked by a high level of plasticity, are activity-dependent structural modifications when some synaptic connections are reinforced and maintained while others are eliminated (Erzurumlu and Gaspar, 2012). This increased level of remodeling in response to environmental stimuli is attenuated in the mature brain when the critical period ends, except for adult neurogenic regions such as the olfactory bulb (OB) and dentate gyrus, which receive new neurons throughout the lifespan of animals (Malvaut and Saghatelyan, 2016, Denoth-Lippuner and Jessberger, 2021). The OB is the first relay of olfactory information processing and is an excellent model for studying how various types of structural modifications are triggered in response to environmental stimuli and learning and how these changes allow for the adaptation of bulbar network functioning and animal behavior.

In the OB, the two main populations of interneurons, granule and periglomerular cells (GCs and PGs), are constantly renewed through the process of adult neurogenesis (Malvaut and Saghatelyan, 2016). GCs establish a particular form of connection with the principal cells of the OB, mitral and tufted cells, through reciprocal dendrodendritic synapses. The inhibition of principal cells by GCs is crucial for olfactory information processing and olfactory behavior (Malvaut and Saghatelyan, 2016). In addition to being renewed, GCs also exhibit a high level of structural plasticity that persists after their full morpho-functional maturation (Hardy and Saghatelyan, 2017). In fact, approximately 20% of fully mature GC dendritic spines are constantly being added/removed from the OB circuitry (Sailor et al., 2016). This persistent structural plasticity optimizes odor information processing in the adult OB (Sailor et al., 2016) and is modulated by odor learning (Wu et al., 2020, Lepousez et al., 2014) and sensory activity (Lledo and Saghatelyan, 2005, Bastien-Dionne et al., 2010). Indeed, it has been shown that odor learning induces an increase in the spine density of GCs (Wu et al., 2020, Lepousez et al., 2014) while sensory deprivation decreases the spine density (Kelsch et al., 2009, Lledo and Saghatelyan, 2005, Bastien-Dionne et al., 2010). Moreover, *in vivo* imaging studies have revealed that another type of structural plasticity involving the relocation of mature spines allows for the rapid adaptation of the OB circuitry to changes in the olfactory environment (Hardy and Saghatelyan, 2017, Breton-Provencher et al., 2016, Sailor et al., 2016). This process has been shown to be specific to mature adult-born GCs (Malvaut et al., 2017, Breton-Provencher et al., 2016, Sailor et al., 2016), which are known to play a unique role in OB functioning compared to their early-born counterparts. Notably, during their integration into the OB neuronal network, two-to three-week-old adult-born OB neurons go through a phase when they are more responsive to new odorants and have broader responsiveness to individual odorants (Belnoue et al., 2011, Magavi et al., 2005). Adult-born GCs also play an important role in a wide range of social and spontaneous olfactory behaviors as well as odor learning and memory (Malvaut and Saghatelyan, 2020). Specifically, these cells play a major role in learning an odor discrimination task (Alonso et al., 2012, Li et al., 2018, Grelat et al., 2018), a task that also induces an increase in the spine density of this population of cells (Wu et al., 2020, Lepousez et al., 2014). Although all these studies highlight changes in spine density, it remains unclear whether the morphometric parameters of spines also change with the level of olfactory stimuli and/or learning. This is of particular relevance given that the size of a spine head is correlated with the surface of the postsynaptic density as well as synaptic strength, whereas both the length and width of a spine neck are linked to postsynaptic potential (Kharazia and Weinberg, 1999, Takumi et al., 1999, Ganeshina et al., 2004, Arellano et al., 2007, Tønnesen et al., 2014).

Historically, dendritic spines have been classified into a few groups, such as stubby, mushroom, or thin, based on their morphological features, which helps simplify data interpretation (Rodriguez et al., 2008, Ghani et al., 2017). However, clustering, an unsupervised learning technique, has emerged as a promising alternative for the analysis of dendritic spines. Clustering groups of spines based on inherent similarities without predefined categories can reveal unknown spine subtypes of functional or pathological significance, capture continuous variations in spine morphology, and reduce subjectivity and bias in classification (Bokota et al., 2016, Pchitskaya and Bezprozvanny, 2020, Ghani et al., 2017). By providing a more nuanced and objective perspective on spine diversity, clustering can enhance our understanding of neuronal function and dysfunction and capture how the morphometric properties of spines are affected by sensory inputs and/or learning and memory. To assess how sensory activity and odor learning of different complexities affect the structural plasticity of adult-born GCs in the OB, we developed a computational pipeline to extract dendritic spines from confocal images and perform cluster analyses of reconstructed dendritic spines based on distinct morphometric features. We discovered an unexpected diversity of structural plasticity of adult-born GCs in response to the level of sensory activities and odor learning of different complexities. Sensory deprivation decreased spine density without any changes in the morphometric properties of the remaining spines or in spine clusters. Interestingly, while complex odor learning increased spine density without any substantial modifications in the morphometric properties of spines, simple odor learning tasks, on the other hand, did not affect spine density but led to the enlargement of remaining spines. Our data reveal that a distinct mode of structural plasticity might be engaged in adult-born GCs to adapt the functioning of the bulbar network in response to the level of sensory input and complexity of odor learning paradigms.

## Results

### Go/no-go odor discrimination learning tasks of different complexities lead to distinct changes in the spine density of adult-born GCs

We first investigated the impact of odor learning tasks of different complexity on the structural plasticity of adult-born GCs. Approximately 4 weeks following an injection of a GFP-encoding lentivirus into the rostral migratory stream (RMS) (**Figure 1A)**, mice were trained on a go/no-go olfactory discrimination task and depending on the complexity of the task, different mixes of odors were presented (**Figure 1B**). In the simple version of the task, the mice had to discriminate between two dissimilar odorants: 1% octanal as S+ vs. 1% decanal as S-. Animals from the learner group (n=4 mice) received a water reward if the odor was discriminated correctly. Animals from the control group (n=3 mice) were exposed to the same conditions but received a reward regardless of the odor. The learner group reached the criterion of 80% of correct responses in 5 blocks over the course of 1 day (**Figure 1C**) and after only 3 blocks, the success rate of the learner group was significantly different from that of the control group (Student t-test, *p*<0.001). Following the learning test, the mice were perfused, and high-resolution confocal images of adult-born GCs were acquired (**Figure 1D**). We first quantified the spine density on the distal dendrites of adult-born GCs in the external plexiform layer, the site of contact between GCs and bulbar principal neurons. Our quantification did not reveal any significant difference in spine density between the control and go/no-go simple learning groups (0.55±0.21 spines/µm for the control group, n=110 cells from 3 mice vs. 0.55±0.17 spines/µm for the simple learner group, n=111 cells from 3 mice, **Figure 1E**).

**Figure 1.**
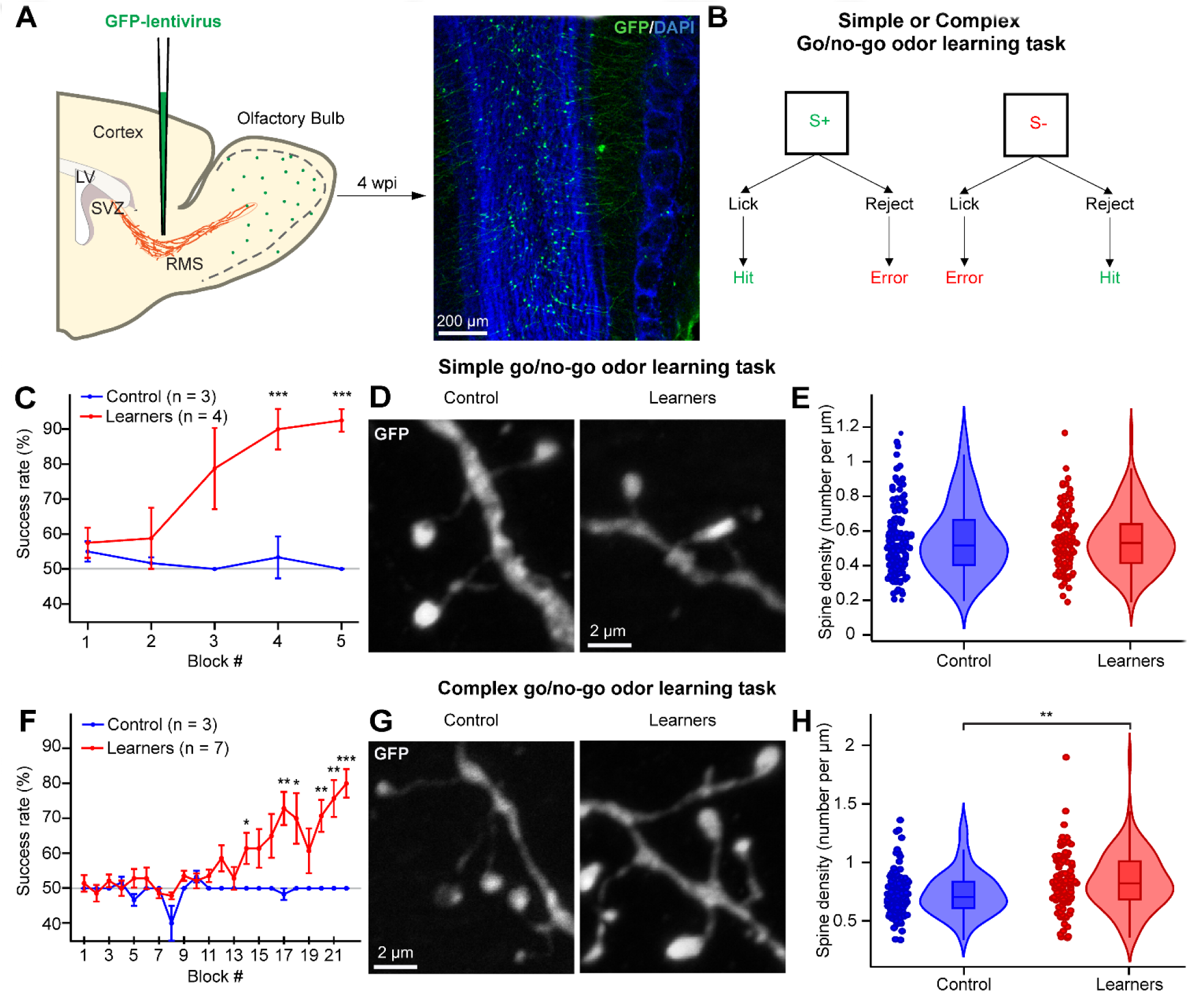
Go/no-go odor discrimination affects the spine density of adult-born GCs differently, depending on the complexity of the task. **(A)** Schematics of the lentiviral injection in the RMS (left) and a low magnification image of the OB (right) showing GFP+ adult-born GCs (green). The OB section was counterstained with DAPI (blue). **(B)** The go/no-odor discrimination task started 4 weeks post-injection. **(C)** Success rate (as a % of total responses) per block for the simple go/no-go task between the control (blue, n=3 mice) and learner (red, n=4 mice) groups. **(D)** Examples of dendritic spines of adult-born GCs in the control and simple go/no-go learner groups. **(E)** Following the simple go/no-go task, there were no significant differences in spine density between cells from the control (blue, n=110 cells from 3 mice) and the learner (red, n=111 cells from 3 mice) groups. **(F)** Percentage of success per block for the complex go/no-go task between the control (blue, n=3 mice) and the learner (red, n=7 mice) groups. **(G)** Examples of dendritic spines of adult-born GCs in the control and complex go/no-go learner groups. **(H)** The complex go/no-go task resulted in a significantly higher spine density in the learner group (red, n=104 cells from 4 mice) compared to the control group (blue, n=97 cells from 4 mice).

Another group of animals was trained with a complex version of the go/no-go task involving very similar odor mixtures: 0.6% limonene(+) and 0.4% limonene(-) as S+ vs. 0.4% limonene(+) and 0.6% limonene(-) as S-. The learner group (n=7 mice) reached the criterion of 80% of correct responses after 22 blocks, while the control group, as expected, never reached this criterion. The difference between the learner and control groups in terms of success rate was highly significant after block 13 (**Figure 1F**). Interestingly, the assessment of the spine density of adult-born GCs revealed that, unlike the simple discrimination task, the complex task led to an increase in spine density after training (control group: 0.73±0.19 spines/µm, n=97 cells from 4 mice vs. learner group: 0.84±0.25 spines/µm, n=104 cells from 4 mice, *p*=0.003 with the Student t-test, **Figure 1G-H**). These data suggest that the spine density of GCs is differently affected depending on the complexity of the go/no-go learning task.

We also observed that the spine density of GCs from mice of a control group undergoing the complex go/no-go task was higher than the spine density of cells from the control group undergoing the go/no-go simple task (0.55±0.21 spines/µm vs. 0.73±0.19 spines/µm for the simple and complex go/no-go control groups, respectively, *p*<0.001with the Student t-test) **(Figures 1E** and **1H)**. As the mice in the complex go/no-go control group were exposed to odors in an operant conditioning task for a much longer period of time and encountered odors of a more complex nature, these data suggest that the duration of olfactory stimulation and the complexity of the odors used might affect the spine density of adult-born GCs. These results are in line with previous observations showing that a lack of odor stimulation decreases the spine density of adult-born GCs (Zuo et al., 2005, Breton-Provencher et al., 2016).

### Odor deprivation decreases the spine density of adult-born GCs

We next investigated the impact of decreased levels of olfactory stimulation on the structural plasticity of adult-born GCs. To do so, 14 days following the injection of a GFP-expressing lentivirus into the RMS, we performed a unilateral nostril occlusion to deprive an OB hemisphere of odor stimulation. We used the contralateral OB hemisphere as a control. Following 14 days of sensory deprivation, we performed a quantitative assessment of the spine density of adult-born GCs. To ensure the effectiveness of the sensory deprivation, we assessed the level of tyrosine hydroxylase (TH) expression, which is known to be regulated in an activity-dependent manner (Bastien-Dionne et al., 2010), in the glomerular layer of the odor-deprived OB hemisphere compared to the control. We observed a characteristic 60% decrease in TH expression in the odor-deprived OB hemisphere (**Figure 2A**). Moreover, and in accordance with previous reports (Zuo et al., 2005, Breton-Provencher et al., 2016), sensory deprivation resulted in a statistically significant decrease in the spine density of adult-born GCs in the odor-deprived OB hemisphere compared to the control OB hemisphere (control OB: 0.55±0.19 spines/µm, n=52 cells from 3 mice vs. occluded OB: 0.44±0.12 spines/µm, n=50 cells from 3 mice, *p*<0.001 with a paired Student t-test; **Figure 2B**-**C**). Our results showed that the level of sensory activity in the OB influenced the plasticity of adult-born GCs by modulating their spine density.

**Figure 2.**
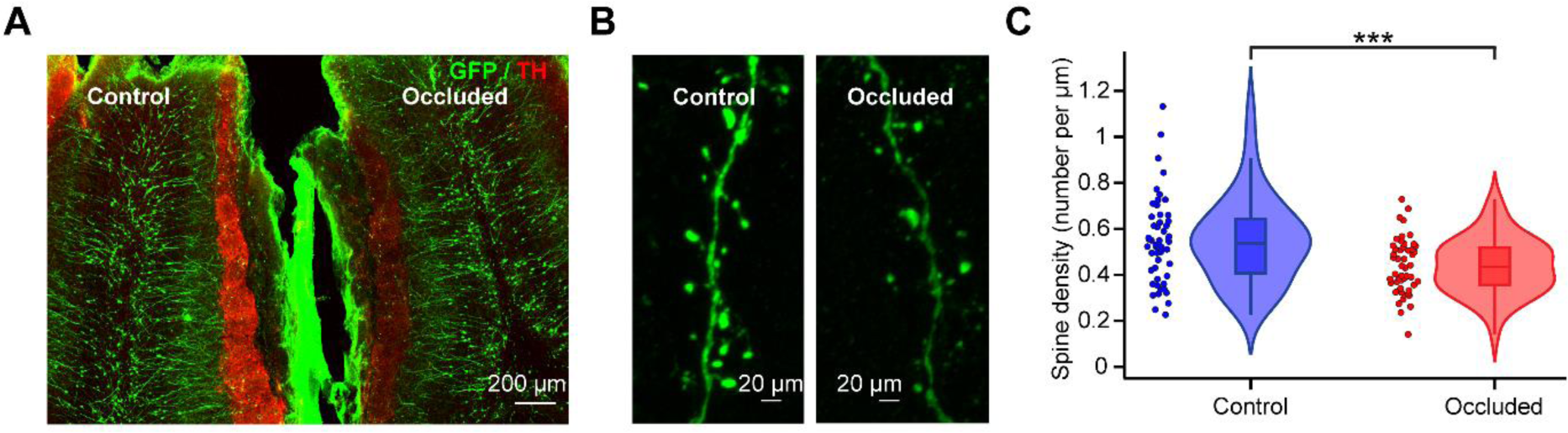
Odor deprivation affects the spine density of adult-born GCs. **(A)** Confocal image showing two OB hemispheres from an unilateral odor-deprived mouse in which 4-week-old adult-born GCs express GFP (green). Following TH labelling (red), a significant decrease in TH labelling was observed in the occluded OB hemisphere. **(B)** Representative confocal images of GFP-labelled secondary dendrites of adult-born GCs in both the control and occluded OB hemispheres. **(C)** Violin plot representation highlighting the significant decrease in the spine density of adult-born GCs following odor deprivation compared to GCs in the control OB (control OB: blue, n=52 cells from 3 mice and odor-deprived OB: red, n=50 cells from 3 mice; *p*<0.001 with a paired Student t-test.

### A pipeline to perform quantitative analyses of 3D-reconstructed dendritic spines from confocal images

Our results show that the spine density of adult-born GCs could be differently affected by odor learning tasks of different complexity and sensory stimulation. However, sensory stimulation and learning may induce changes not only in the number of spines but may also affect the morphometric properties of existing spines. We thus developed a pipeline that starts with the 3D-reconstruction of spines, followed by an assessment of up to 10 distinct morphometric parameters, dimension reduction, and cluster analysis. High resolution confocal images of dendritic spines from animals from the different experimental conditions were first obtained (**Figure 3A**). The images were deconvolved using the DAMAS algorithm (Brooks and Humphreys, 2006, Dougherty, 2005), followed by segmentation with the morphological snake Chan-Vese algorithm (Viola and Wells, 1995, Alvarez et al., 2010, Chan and Vese, 2001, Marquez-Neila et al., 2013). Lastly, a marching cube algorithm was used to reconstruct the mesh (Lewiner et al., 2003, Lorensen and Cline, 1987). We extracted 4936 spines using the selection tool of Meshlab. For each extracted spine, 10 features were calculated: Length, Surface, Volume, Average Distance, Open Angle, Hull Volume, Hull Ratio, Coefficient of Variation in Distance (CVD), Mean Curvature, and Mean Gaussian Curvature (**Figure 3B**). A covariance analysis was performed to find the correlation between the features and to only keep the least correlated ones (**Table 1**). Correlations ranging from 0.7 to 0.89 were considered strong, while correlations ranging from 0.9 to 1 were considered very strong based on the literature (Schober et al., 2018). A very strong correlation was discovered between Length and Average Distance (0.97). For further analyses, we selected the Length parameter because it is a widely accepted and used feature. The Surface, Volume, and Hull Volume parameters also showed very strong correlations (0.98 between Surface and Volume, 0.97 between Surface and Hull Volume, and 0.99 between Volume and Hull Volume). Of those three parameters, we selected Surface as it is a widely used feature in morphometric assessments of dendritic spines. Lastly, the Hull Ratio, CVD, and Open Angle were retained because they were not correlated with any other feature. We next applied a Standard Scaler followed by a dimension reduction with a Principal Component Analysis (PCA) (**Figure 3C**). This led to three Principal Components (PCs) representing 85.4% of the total variance of the data. Spines from the three experiments were included (go/no-go simple task, go/no-go complex task, and sensory deprivation), together with their respective controls. The resulting representation of the dendritic spines was homogenous, with no obvious clusters (**Figures 3D, 3E,** and **3F**). Based on the analysis of the directionality of the coefficients of the linear combination (PCA loadings), PC1 was mostly composed of Surface, Length, and CVD, PC2 was composed of Surface and Open Angle, and, lastly, PC3 was mostly related to the Hull Ratio. The extracted features could then be used for further analyses such as clustering.

**Figure 3.**
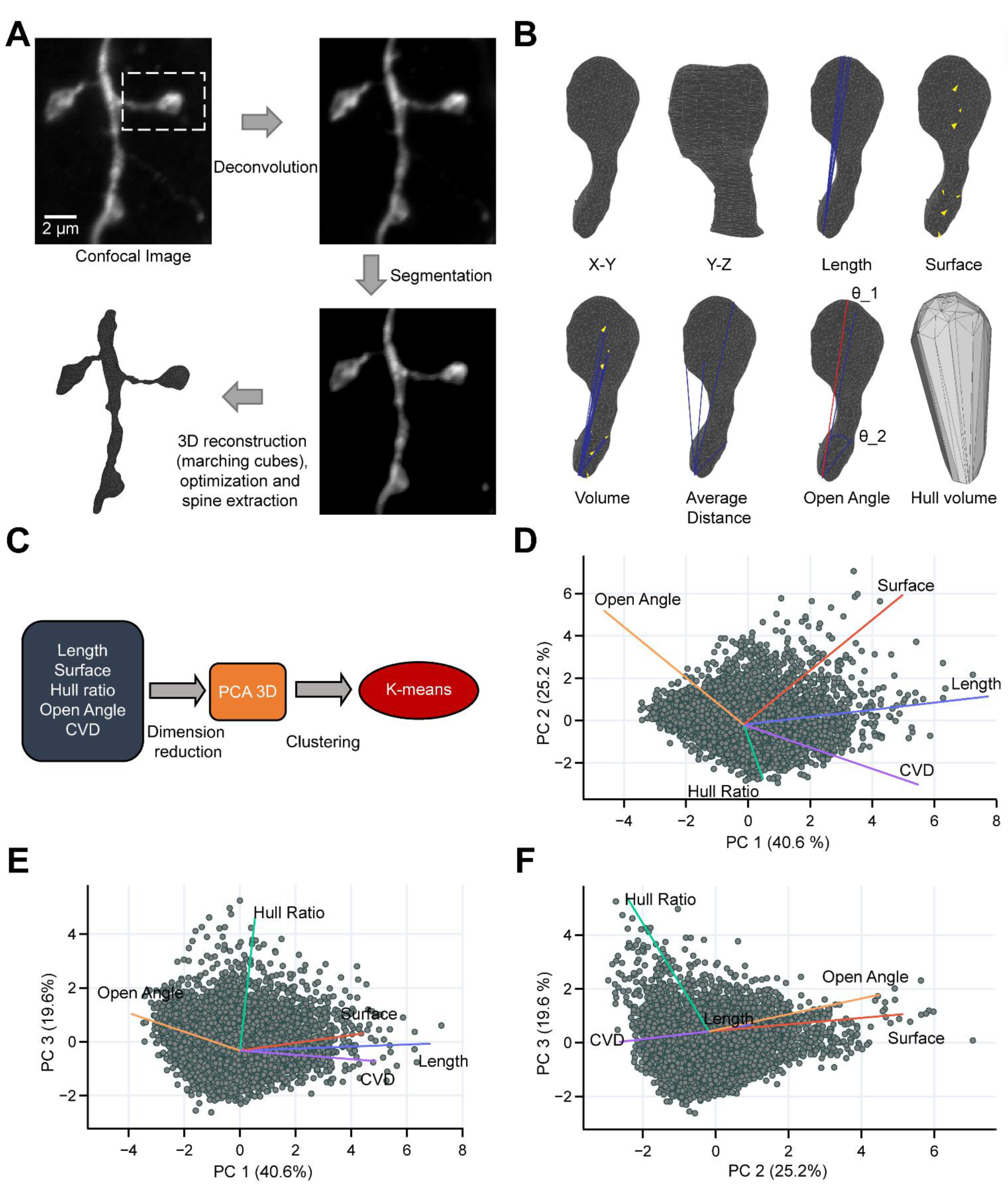
Reconstruction of dendritic spines from confocal microscopy images and analysis pipeline. **(A)** Confocal image (top left) of a dendritic segment of an adult-born GC where the spine of interest is highlighted by a dashed rectangle. The same image after deconvolution (top right), segmentation (bottom right), and reconstruction (bottom left). **(B)** Different representations of the dendritic spine of interest reconstructed in 3D with calculated morphometric features. **(C**) General pipeline to analyze dendritic spines based on a combination of dimension reduction and statistical methods. **(D-F)** Biplot representations of all reconstructed spines after PCA, where each grey point represents a dendritic spine. Each line represents the directionality of the coefficients of the linear combination (PCA loadings) for each feature.

**Table 1.**
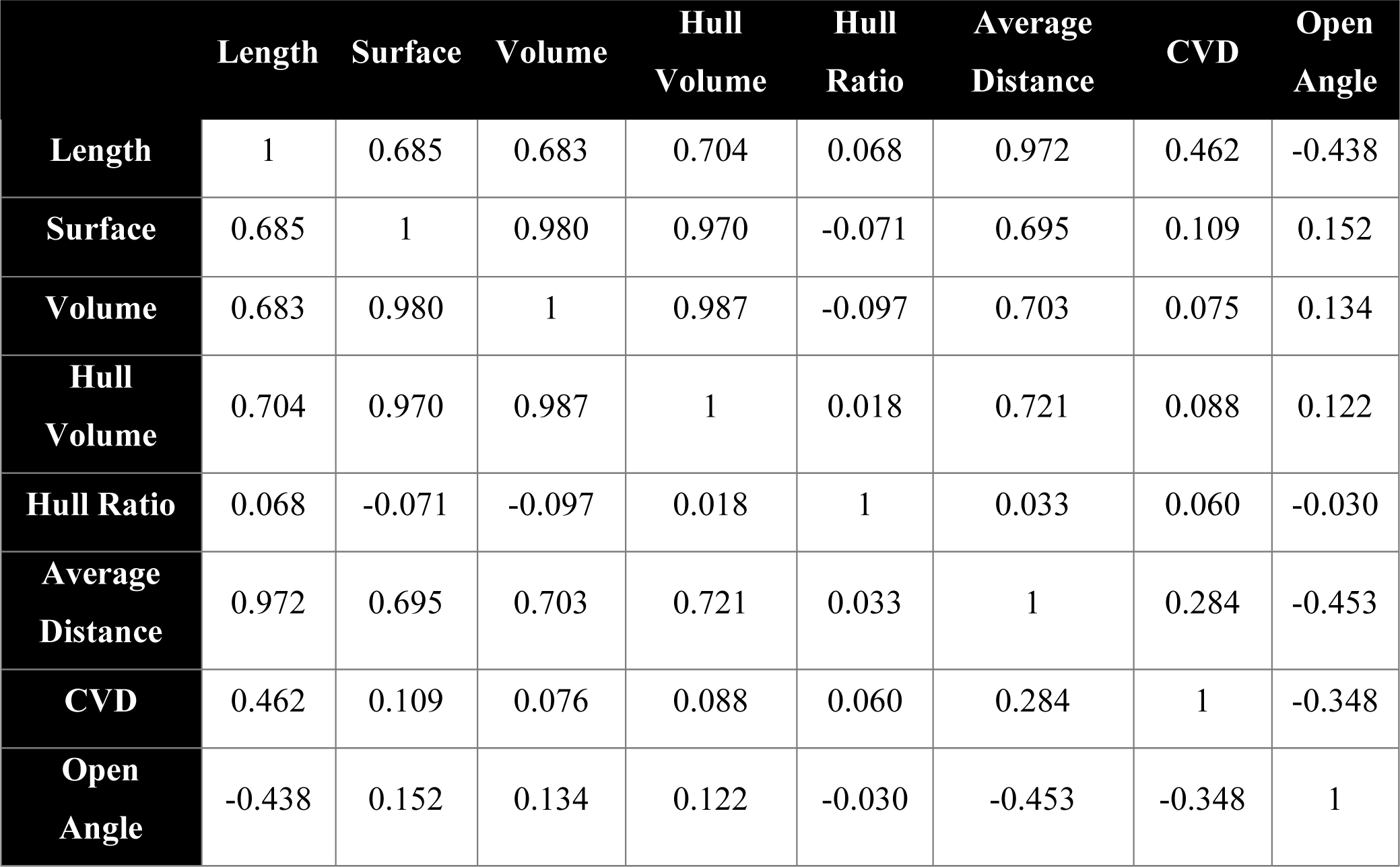
Pearson’s correlation coefficients between the morphometric features: Length, Surface, Volume, Hull Volume, Hull Ratio, Average Distance, CVD, and Open Angle

### Adult-born GC spines can be assembled into different clusters based on experimental conditions

Dendritic spines are highly dynamic, and these protrusions constantly change in size and shape in response to the sensory environment. As there is a continuum of spine shapes and sizes (Yuste and Bonhoeffer, 2004), which can also be seen in our dataset after dimension reduction, any attempt to classify spines based on their shape using arbitrary categories could be erroneous (Pchitskaya and Bezprozvanny, 2020). As such, an automatic clustering approach is preferable. In such an approach, the number of clusters is the first parameter that needs to be determined. We identified the appropriate number of clusters for our spine dataset using three different scores: the Elbow score, the Silhouette score, and the Calinski-Harabsz score (**Figure 4A and Figure S1**). All three scores suggested that five clusters were present in the dataset: Cluster 1 (n=1160 spines) was located on the low value of the first principal component (mean PC1=–1.683). Cluster 2 (n=790 spines) was located on the high value of the first principal component (mean PC1=2.146). Cluster 3 (n=628 spines) was located in the middle of the first principal component (mean PC1=0.551), but with a high PC2 value (mean=1.866). Cluster 4 (n=1610 spines) was located at the low value of PC3 (mean=–0.860) and cluster 5 (n=748 spines) was located at the high value of PC3 (mean=1.095) and the low value of PC2 (mean=–1.224) (**Figure 4B**). An assessment of the reconstructed spines in each cluster made it possible to associate the mean value of each feature to the morphology of the spine (**Figure 4C**). Cluster 1 was mainly composed of small stubby spines or small mushrooms, with a short length (1.39±0.31 μm), small surface area (8.24±4.03 μm^2^), small CVD (0.28±0.08), and particularly high Open Angle (0.94±0.11). Cluster 2 was composed of long mushrooms with a long length (3.59±0.81 μm), high CVD (0.45±0.07), and very high surface area (19.35±6.65 μm^2^). Cluster 3 was composed of spines with a large surface area (26.17±9.18 μm^2^). However, these spines were generally shorter than the spines in cluster 2 (2.55±0.51 μm) and had a higher Open Angle (1.01±0.16). Clusters 4 and 5 had similar spines, with similar lengths (1.89±0.31 μm and 2.07±0.52 μm, respectively), similar surface areas (9.36±4.16 and 8.42±3.88, respectively), similar Open Angles (0.69±0.16 and 0.66±0.17, respectively), and similar CVDs (0.41±0.05 and 0.451±0.05, respectively). However, the spines of cluster 5 were more elongated with higher Hull Ratio (0.85±0.20 vs. 0.40±0.11 for clusters 5 and 4, respectively) which, for these spines, reflect a lower volume as compared to spines of cluster 4. The clustering of dendritic spines of adult-born GCs based on their morphometric features allowed us to determine whether these properties were differently affected in response to a simple or complex go/no-go odor learning task and to sensory deprivation.

**Figure 4.**
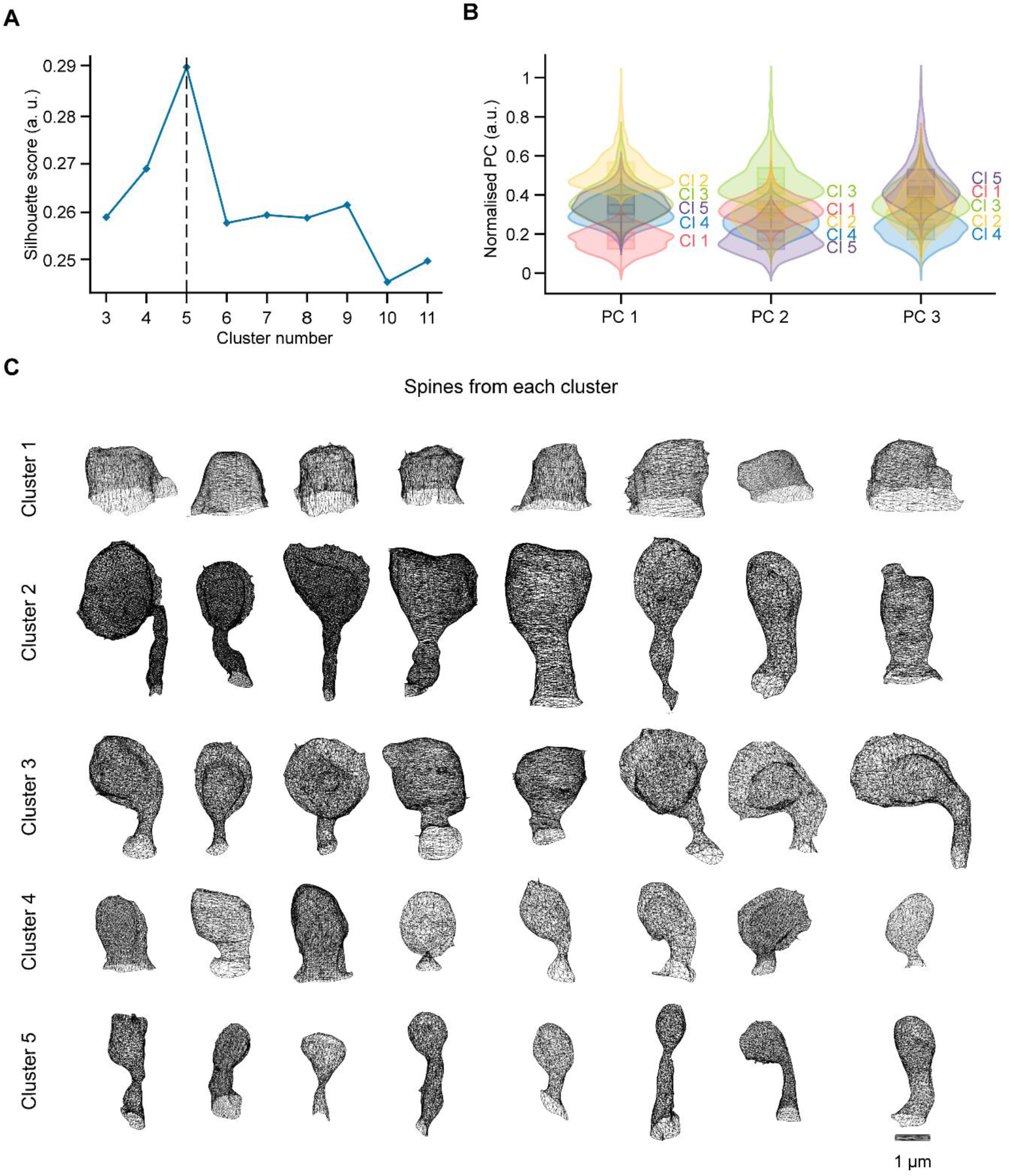
Clustering of spines of adult-born GCs based on their morphological features. (**A**) Representation of the silhouette score for 3 to 11 clusters. The dashed line shows the optimal number of clusters. (**B**) Violin plot representation of each principal component (PC) after normalization for five clusters. (**C**) Morphological representation of 10 reconstructed spines for each of the five clusters taken near the center of each cluster.

**Figure S1.**
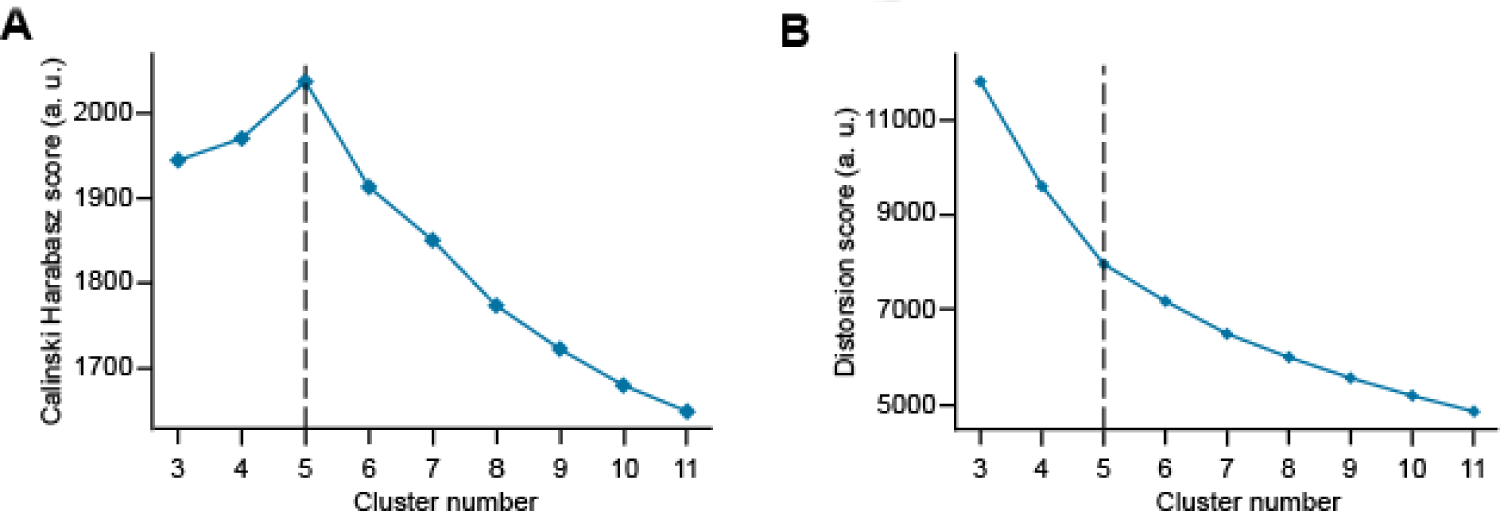
***The different clustering scores also suggested that our spine dataset could be divided into five distinct clusters.*** ***(A)** The Calinski-Harabsz score and **(B)** the Elbow score showed that the most likely number of clusters was 5.*

### Sensory deprivation does not affect spine morphology

Our results, as well as those from previous reports (Zuo et al., 2005, Breton-Provencher et al., 2016), indicate that sensory deprivation led to a decrease in the spine density of adult-born GCs. It remains unknown, however, whether sensory deprivation also changes the morphometric properties of the remaining spines. To address this issue, we compared the morphometric features of GC spines in the occluded odor-deprived group (n=370 spines) to those of GC spines in the control group (n=457 spines). We observed no significant differences in any of the morphometric properties of GC spines between the odor-deprived and control groups (**Figure 5A**). Cohen’s term d (effect size index) showed a very small to small effect for every variable. After clustering, no differences in cluster distribution were seen with the Agresti-Caffo test (**Figure 5B**). The most prevalent clusters of spines were clusters 4 and 5, which contained similar spines. These spines were relatively small with thin-like spines that together represented 50.4% and 56% of the total spines for the control and occluded groups, respectively. Spines with a high surface area (clusters 2 and 3) had a representation of 31.5% and 28.4% for the control and occluded groups, respectively. To ensure that there were no changes in morphologies within the cluster, we compared the occluded and control groups for each cluster. No differences were seen in spine morphologies for any cluster (**Figure 5C**). This result suggests that sensory deprivation via nostril occlusion only affected the spine density of adult-born GCs, with no changes in the morphometric properties of the remaining spines.

**Figure 5.**
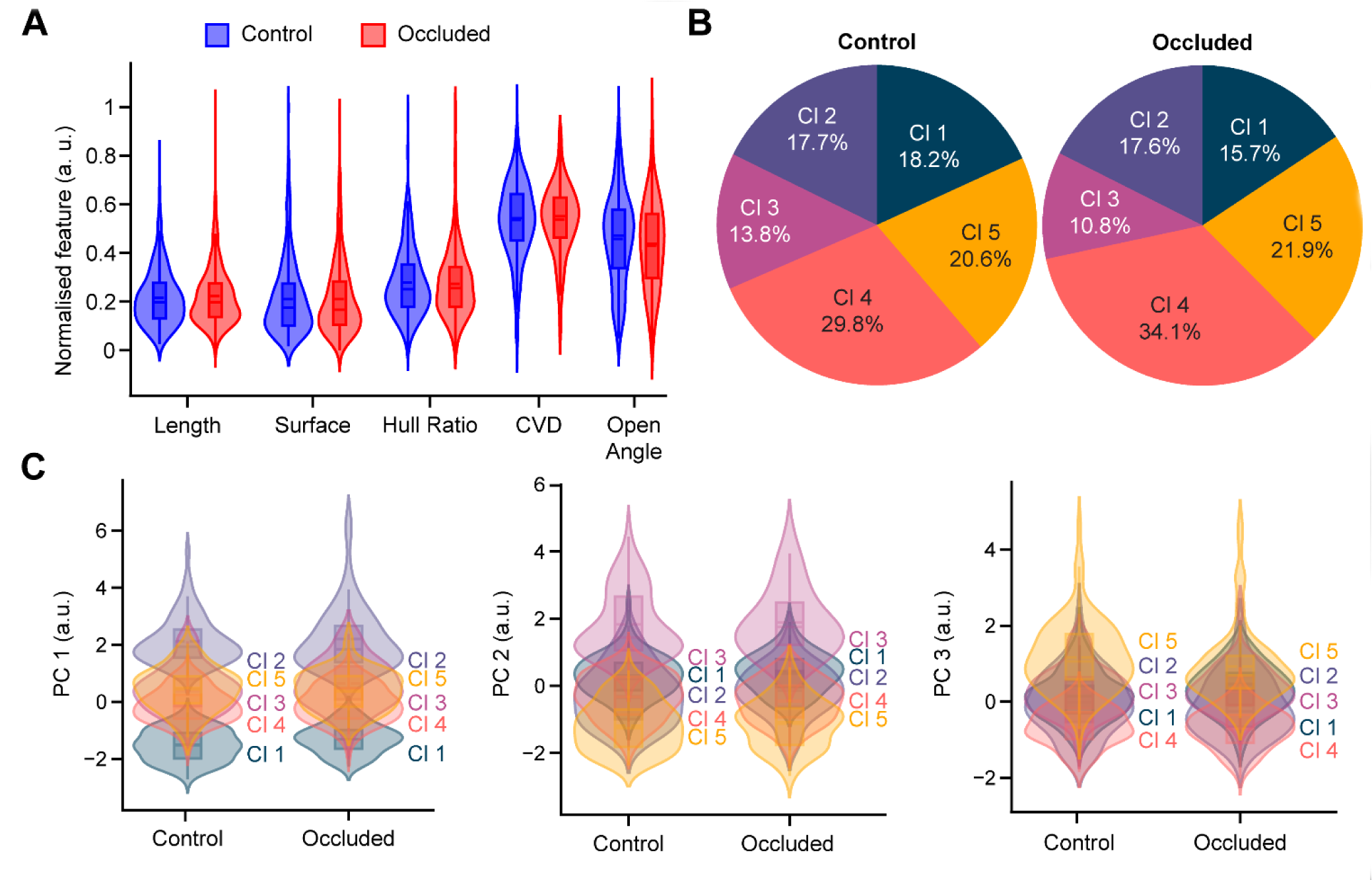
Sensory deprivation does not affect dendritic spine morphologies. (**A**) Violin plot representation of spines from the control (blue) and occluded (red) groups for each normalized feature. (**B**) Pie chart representation of the cluster distributions in the control and occluded groups. (**C**) Violin plot representation of each principal component after dimension reduction and normalization for five clusters for the control and occluded groups.

### Different types of plasticity are evoked by learning, depending on the complexity of the go/no-go task

Given the distinct effect on spine density of adult-born GCs in the control groups of go/no-go simple and complex odor learning tasks, we determined whether the complexity of the operant task also influenced the morphology of dendritic spines. We compared spines from both the simple (n=1334 spines) and complex task (n=832 spines) control groups. Interestingly, we observed major differences between spines from both control groups. Cohen’s term’ size effect test reported a very small to small effect for every variable. When both control groups were compared, significant differences were found with respect to spine surfaces (13.37±8.90 μm^2^ vs. 10.95±7.16 μm^2^ for the simple and complex task control groups, respectively; *p*<0.001 with the Student t-test), Hull Ratio (simple task control group: 0.60±0.21 vs. 0.50±0.19 for the simple and complex task control groups, respectively; p<0.001 with the Student t-test), CVD (0.41±0.09 vs. 0.36±0.09 for the simple and complex task control groups, respectively; *p*<0.001 with the Student t-test), and Open Angle (0.85±0.19 vs. 0.64±0.19 for the simple and complex task control groups, respectively; *p*<0.001 with the Student t-test, **Figure S2A**). No difference was observed with respect to spine length. The cluster distribution between the two control groups was also different. There were significantly more spines with cluster 2 (Agresti-Caffo, *p*<0.001) and cluster 4 (Agresti-Caffo, *p*<0.001) in the control group of the complex go/no-go task, which concomitantly resulted in fewer spines with a cluster 3 morphology (Agresti-Caffo, *p*<0.001) and a cluster 5 morphology (Agresti-Caffo, *p*<0.001) (**Figure S2B**). The duration of the odor stimulation in the go/no-go task and the complexity of the odor used thus had an impact on the morphology of the dendritic spines, even when no association between a particular odorant and a reward was involved.

**Figure S2.**
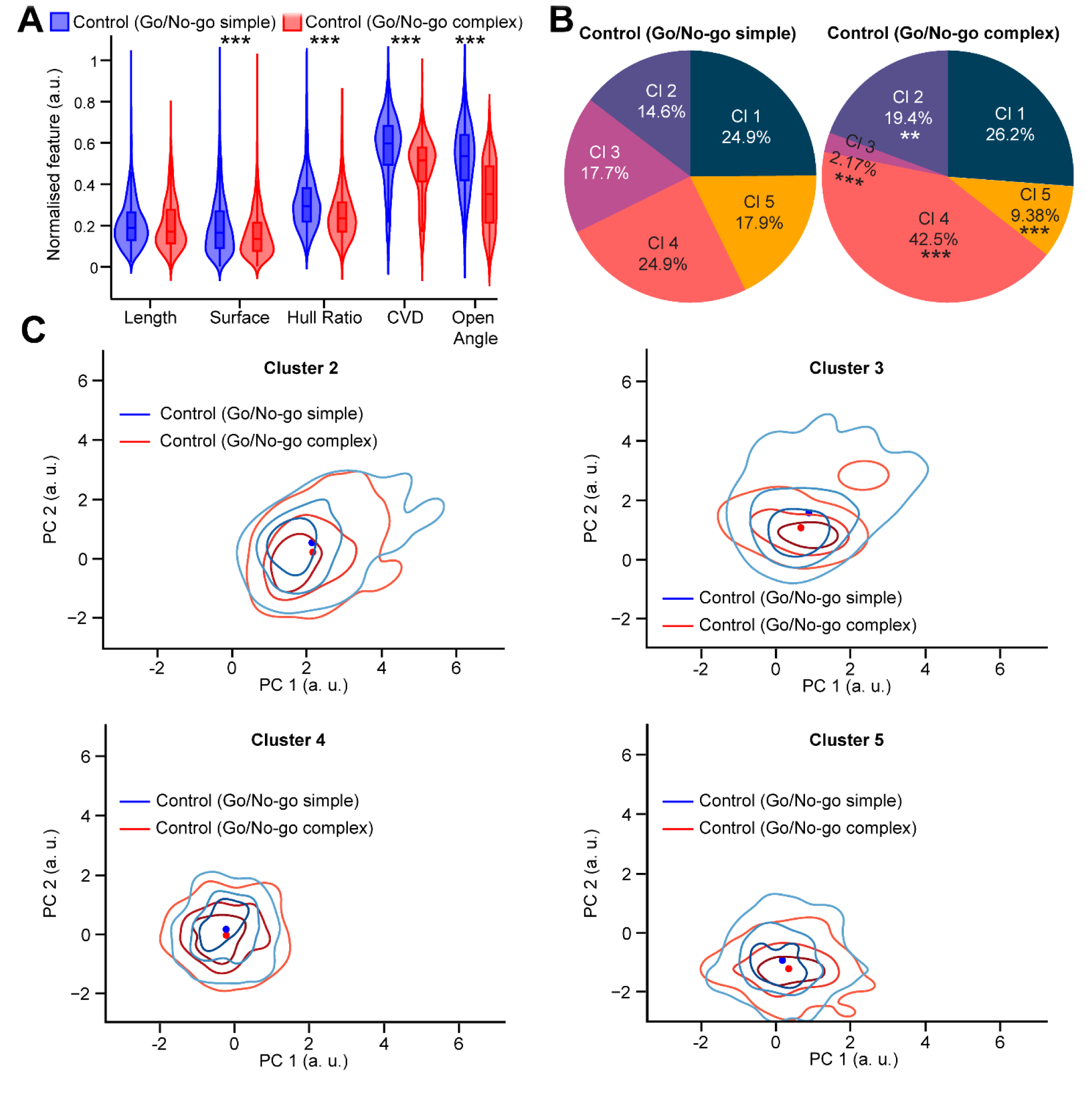
Major morphological differences exist between the control groups of the simple and complex go/no-go tasks. (**A**) Violin plot representation of the control group of the simple task in blue and the control group of the complex task in red for Length, Surface, Hull Ratio, CVD, and Open Angle. (**B**) Pie chart representation of the cluster distribution in the control group of the simple task and the control group of the complex task. (C) KDE maps of the representation of the spine densities of clusters 2, 3, 4, and 5 in the space of PC1 and PC2 for the control group of the simple task in blue and the control of the complex task in red.

We next determined whether learning a go/no-go simple task that did not induce any changes in the spine density of adult-born GCs affected the morphometric properties of spines. The morphometric properties of spines from the learner group of the simple go/no-go task (n=767 spines from 111 cells, 3 mice) were compared to those of the control group (n=1334 spines from 119 cells, 3 mice). Cohen’s term d was calculated and indicated a very small to small effect for every variable. Significant differences (**Figure 6A**) were found for four features between the control and learner groups: Surface, with 13.37±9.0 μm^2^ and 16.19±10.40 μm^2^ (*p*<0.001 with the Student t-test) for the control and learner groups, respectively, Hull Ratio, with 0.6±0.21 and 0.57±0.20 (*p*<0.05 with the Student t-test) for the control and learner groups, respectively, CVD, with 0.41±0.09 and 0.4±0.09 (*p*<0.01 with the Student t-test) for the control and learner groups, respectively, and Open Angle, with 0.85±0.19 and 0.93±0.18 (*p*<0.001 with the Student t-test) for the control and learner groups, respectively. No significant changes were observed in spine length between the two groups. These results indicate that while spine length was not affected by the simple go/no-go learning task, this learning paradigm led to larger spines, as attested to by changes in the Surface, Hull Ratio, CVD, and Open Angle.

**Figure 6.**
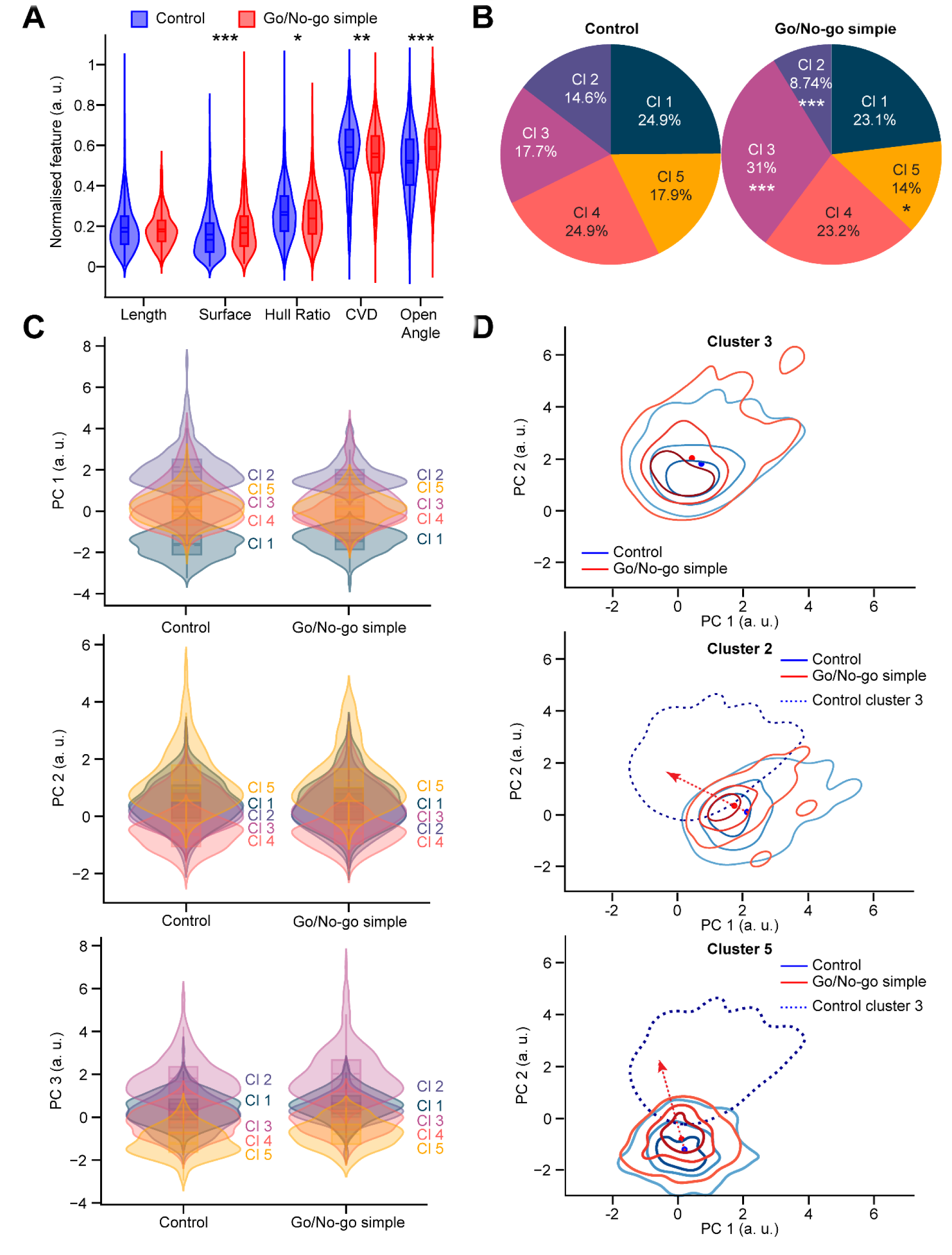
A simple go/no-go task leads to changes in spine morphology. (**A**) Violin plot representation of each normalized morphometric feature for the control (blue) and learner (red) groups. **(B)** Pie chart representation of the cluster distribution in the control and go/no-go learner groups performing a simple odor discrimination task. Note the decrease in the representation of clusters 2 and 5, as well as the increase in cluster 3 following learning. **(C)** Violin plot representation of the five clusters after dimension reduction and for each normalized PC for the control and learner groups. **(D)** KDE maps of the representation of the spine densities of clusters 3, 2, and 5 in the space of PCs 1 and 2 for the control group in blue and the learner group in red. The dotted line corresponds to the KDE representation of the spine density of cluster 3. The red arrow represents the direction of the cluster from the center of the KDE map for the control group to the center of the KDE map for the go/no-go simple group.

The changes in morphology were also significant with respect to the distribution of several clusters (**Figure 6B**). Following learning, the proportion of spines in cluster 2 decreased from 14.6% in the control group to 8.74% in the learner group (Agresti-Caffo, *p*<0.001). Similarly, the spines of cluster 5 decreased from 17.9% in the control group to 14% in the learner group (Agresti-Caffo, *p*<0.05). The proportion of spines in cluster 3 increased from 17.7% in the control group to 31% in the learner group (Agresti-Caffo, *p*<0.001). Overall, learning to associate the simple odor with a water reward in go/no-go odor discrimination task led to the appearance of more spines with a mushroom-like shape as seen by an increase in the Surface and Open Angle and a reduction in the Hull Ratio and CVD. To ascertain that each cluster in both groups were represented by spines of similar morphologies, the spines of each cluster in the control group were compared to those of the learner go/no-go group. No differences were seen in spine morphologies within any cluster (**Figure 6C**). To understand the morphological changes between clusters, KDE was used to map the spine densities of clusters 2, 5, and 3 in the space of PC1 and PC2 (**Figure 6D**). After learning, spines from cluster 2, which are represented by long mushrooms, and spines from cluster 5, which are represented by thin spines with complex structures, tended to move toward cluster 3. These observations thus implied that spines from clusters 2 and 5, but not spines from clusters 1 and 4, tended to change their morphology toward that of spines from cluster 3, i.e., mushroom-like spines with large surfaces.

As complex go/no-go odor discrimination tasks resulted in an increase in spine density (**Figure 2F**), we next determined whether this task also induced changes in spine morphology. The control group (n=832 spines from 99 cells, 4 mice) was compared to the learner group (n=1175 spines from 112 cells, 4 mice) that performed the complex go/no-go task. Interestingly, no significant differences (**Figure 7A**) were found for any of the morphometric features. Cohen’s term d reported a very small to small effect for every variable. We observed only a slight change in the distribution of clusters (**Figure 7B**) with the population of cluster 5 increasing (Agresti-Caffo, *p*<0.05) between the control (9.38%) and the learner (12.7%) groups. The spines in cluster 3, which are mushroom-like spines with large surface areas, were particularly underrepresented, with only 2.17% and 2.55% in the control and learner groups, respectively. However, cluster 4, which contains thin spines with no complex shapes, constituted 42.5% of the control group and 41.1% of the learner group. To ascertain whether the clusters were represented by spines with the same morphologies, we compared the control and learner go/no-go groups for each cluster. No differences were observed in spine morphologies within any cluster (**Figure 7C**). To understand the changes in morphologies, KDE maps of spine densities for clusters 2, 3, and 5 were computed (**Figure 7D)**. The differences in the distribution of spines in cluster 3 and cluster 2 were not significant between the control and learner groups. There were significant differences between the control and learner groups for the spines in cluster 5 that tended to change their morphology in the opposite direction to cluster 3, which could explain the low representation of spines in cluster 3 and the high number of spines with a cluster 5 morphology. This is in contrast to the changes observed in the simple go/no learning task when the spines belonging to cluster 5 tended towards cluster 3.

**Figure 7.**
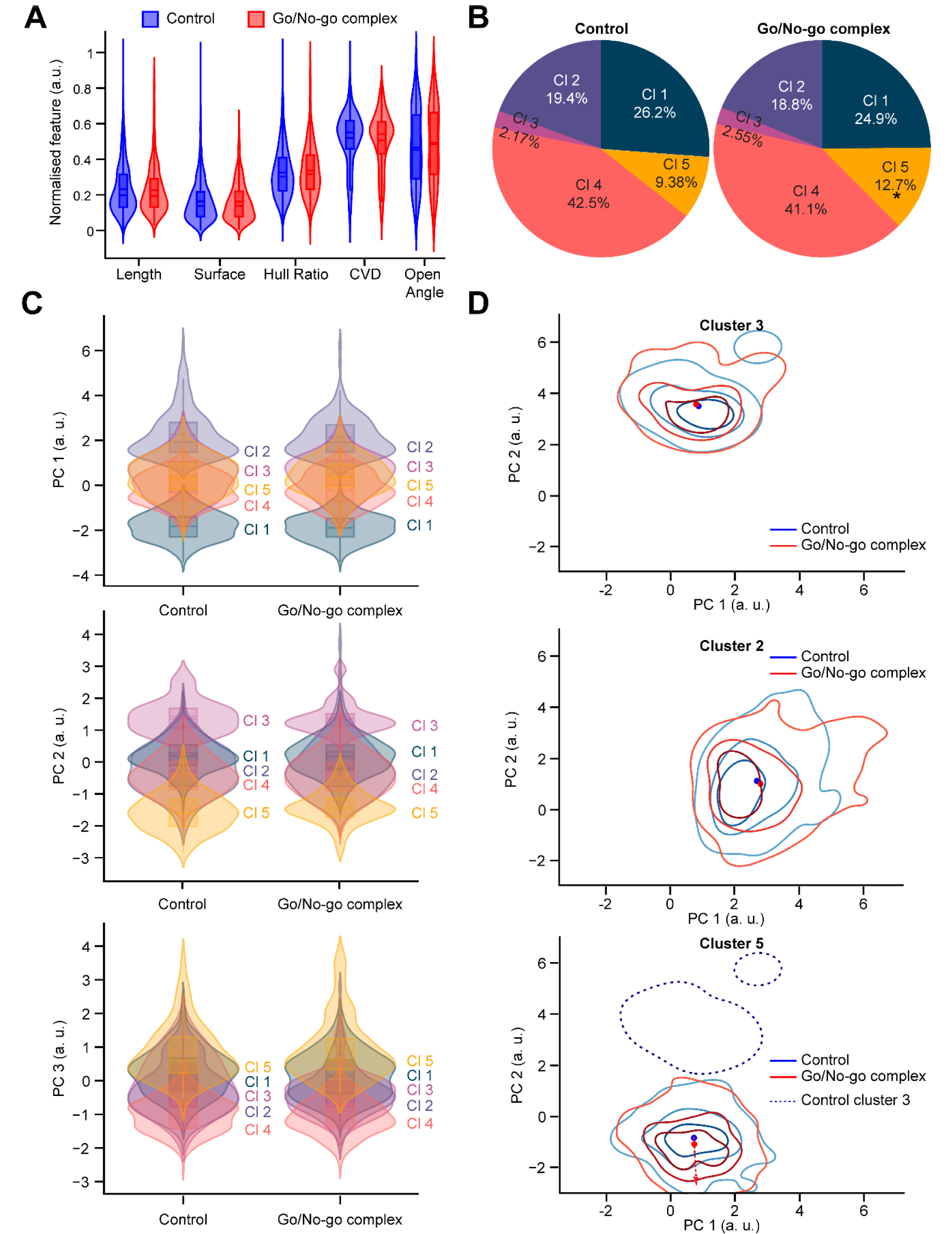
Complex go/no-go odor learning does not lead to changes in spine morphology. (**A**) Violin plot representation of the control group in blue and the learner group in red for Length, Surface, Hull Ratio, CVD, and Open Angle after normalization for the morphometric features. **(B)** Pie chart representation of the cluster distribution in the control and learner groups. **(C)** Violin plot representation of each PC after dimension reduction and normalization for five clusters for the control and learner groups. **(D)** KDE maps of the representation of the spine density for clusters 3, 2, and 5 in the space of PC1 and PC2 for the control group in blue and the learner group in red. The dotted line corresponds to the KDE representation of the spine density of cluster 3. The red arrow represents the direction of the cluster from the center of the KDE map for the control group to the center of the KDE map for the go/no-go simple group.

Overall, our data indicate that, depending on the level of sensory stimulation and the complexity of the odor learning paradigms, distinct forms of structural plasticity of adult-born GCs might be involved in adapting the functioning of the bulbar network. Sensory deprivation decreased spine density, with no changes in spine morphology. The simple go/no-go odor learning task led to changes in the morphometric properties of existing spines without any modifications in spine density. On the other hand, the complex go/no-go learning task increased spine density, with only subtle changes in the morphometric properties of spines. The present study thus shows that there was a vast panoply of distinct forms of structural modifications in the bulbar network.

## Discussion

In the present work we show that the spines of adult-born GCs exhibited different modes of structural plasticity in response to the level of sensory activity and the complexities of odor learning tasks. The computational pipeline we developed enabled us to divide the reconstructed spines into five distinct clusters based on various morphometric properties and to assess how the level of sensory stimulation or odor learning affected the morphology and type of spines. We show that sensory deprivation led to a uniform reduction in spine density, with no changes in the morphology of the spines or their distribution in the different clusters. The difficulty of a go/no-go odor learning task determined whether the structural plasticity of adult-born GCs was reflected by changes in the morphometric properties of the spines or spine density. In fact, while a simple odor discrimination task affected the morphology of the spines, the complex odor learning task resulted in a higher spine density, with no substantial differences in the morphometric properties.

### Distinct types of structural plasticity used by adult-born GCs to adapt the functioning of the bulbar network

Why are such distinct types of structural plasticity of adult-born GCs engaged in response to the level of sensory input and the complexity of odor learning tasks? While further work is required to understand what the functional properties of each spine cluster of adult-born GCs are and how each cluster is involved in odor information processing, it is conceivable that the type of structural plasticity of adult-born GCs is tailored to the needs of the bulbar network in order to adjust its functioning to changing environmental conditions and learned experiences. Sensory deprivation led to an overall reduction in spine density. While these data are consistent with previous reports (Zuo et al., 2005, Breton-Provencher et al., 2016), it was unknown how the morphometric properties of the remaining spines are affected. Our computational pipeline and clustering analysis show that sensory deprivation did not lead to any changes in the morphometric properties of spines and the clusters. GCs synchronize the activities of principal OB neurons through their dendro-dendritic reciprocal synapses, which allows odor information to be processed in the bulbar network (Lagier et al., 2004, Fukunaga et al., 2014). Adult-born neurons may synchronize the activities of new assemblies of principal neurons involved in new memory (Alonso et al., 2012, Sailor et al., 2017, Lledo et al., 2006, Forest et al., 2019). The global reduction in odor-induced activity through unilateral nostril occlusion may not require the synchronization of new assemblies of principal cells because of the lack of sensory inputs. The OB operational neuronal network may thus be maintained through other mechanisms, with no additional structural plasticity required by GC spines. Indeed, it has been shown that sensory deprivation affects not only the output synapses of GCs, as evidenced by the decrease in spine density, but also induces an increase in the input synapses onto GCs (Kelsch et al., 2009). Furthermore, sensory deprivation also triggers an increase in the excitability of newborn GCs, leading to an increase in action potential-dependent GABA release and unaltered synchronized activity of principal cells (Saghatelyan et al., 2005). These studies highlight the adaptive responses in the OB network and indicate that increases in excitability and input synapses onto adult-born GCs maintain the functioning of the neuronal network with no additional need for changes in the morphometric properties of GC spines.

On the other hand, depending on the complexity of the olfactory learning task, the structural plasticity of adult-born GCs is manifested either by changes in the morphometric properties of spines and/or by increases in spine density. The simple go/no-go odor task depended on the presentation of two distinct odors that mice were able to learn very quickly, i.e., in 5 blocks over the course of 1 day (**Figure 1C**). This contrasts with the complex go/no-go learning task based on discrimination between two very similar odor mixtures, which took the mice 22 blocks over multiple days to learn (**Figure 1C**). Our data suggest that learning a simple odor discrimination task might be achieved by enlarging the pre-existing spines of adult-born GCs without forming new ones. In contrast, the complex odor discrimination task required the formation and stabilization of additional spines of adult-born GCs that might be necessary for synchronizing the activities of new principal cell assemblies. These data are in line with previous reports showing that there is an increase in the spine density of adult-born but not pre-existing GCs during a complex odor learning task (Wu et al., 2020, Lepousez et al., 2014). Interestingly, this increase is driven by inputs from the piriform cortex (Wu et al., 2020, Lepousez et al., 2014). Optogenetic stimulation of these projections strengthens learning-induced plasticity (Lepousez et al., 2014), while their inactivation impedes learning-induced changes (Wu et al., 2020). It remains largely unknown how a simple odor learning task affects the feedback projections onto the OB and whether the changes in spine morphologies observed during this task are driven locally in the bulbar network or are mediated by centrifugal projections from other brain regions. It is conceivable that, depending on the complexity of the odor learning task, distinct brain regions or different levels of feedback are required to associate the water reward with odor stimuli and trigger distinct types of structural plasticity. In line with this, it has been shown that different types of odor-associative learning tasks may induce distinct patterns of early immediate gene expression in the orbital and infralimbic cortices (Mandairon et al., 2011), which are known to be involved in odor memory (Tronel and Sara, 2002). Future experiments based on mapping the activated neuronal assemblies across the brain regions during simple and complex go/no-go odor learning tasks will be required to address this issue.

### Spine clustering versus classification: toward the determination of spine clusters with a high level of structural plasticity

The reconstruction of dendritic spines in three dimensions made it possible to obtain more morphometric information than analyses of 2D images of spines. The most relevant features were Length, Surface, Hull Ratio, CVD, and Open Angle, followed by PCA, which reduced the number of dimensions. As spine shapes form a continuum and cannot be categorized (Luengo-Sanchez et al., 2018), the clustering was performed on dendritic spines based on different morphometric properties. When the spines in each cluster were examined, a general characteristic for each cluster could be determined. Cluster 1 predominantly consisted of small stubby spines, cluster 2 of long mushroom spines, cluster 3 of mushroom spines with a larger surface area, and clusters 4 and 5 of long thin spines distinguished by a higher Hull Ratio for cluster 5 (**Figure 4C**). The Open Angle appeared to be a highly effective indicator for distinguishing between mushroom and stubby spines, with the mean value being larger for the stubby spines. The CVD was particularly high for mushroom spines because of their distinct morphology, which is comprised of a head and a neck. The Hull Ratio represents elongated spines with a lower volume. Although all these parameters allowed us to cluster spines based on their morphometric properties, our KDE map representation of spine clusters also made it possible to highlight clusters that were particularly sensitive to odor learning and, at the same time, determine the transition of clusters from one to another within their continuum. Spines in cluster 1 were particularly stable and did not undergo any modifications in response to sensory deprivation or to a simple or complex odor learning task. This contrasts with spines in the other clusters that were modified by learning to different degrees. Although the simple odor learning task induced changes in clusters 2, 3, and 5, the KDE map representation shows that spines in clusters 2 and 5 globally changed their morphology toward the morphology of spines in cluster 3. Clusters 2 and 5 consisted of mushroom spines with smaller surface areas and long thin spines, respectively, whereas cluster 3 harbored mushroom spines with a larger surface area. The transition of clusters 2 and 5 to cluster 3 indicated that these spines became bigger in response to a simple go/no-go odor learning task. The larger spines are usually associated with a larger postsynaptic density (PSD), which might lead to increased synaptic efficacy. It should be mentioned, however, that the spines of GCs are constituents of dendro-dendritic reciprocal synapses, which implies that the same spine contains post-synaptic glutamatergic and pre-synaptic GABAergic parts. It is currently unclear whether the increased surface area reflects changes in both the pre-and post-synaptic constituents and what functional outcome is induced by these changes.

Interestingly, while the complex odor learning task largely resulted in changes in spine density, the subtle morphometric modifications observed after this learning task were opposite to those of the simple go/no-go task. Our cluster analysis and KDE map representation show that there was an increase in spines in cluster 5 while the spines in the cluster 3 were particularly under-represented in both the control (2.17%) and learner (2.55%) groups. The spines in cluster 5 tended to change their morphology in a direction opposite to cluster 3, unlike with the simple go/no-go learning task where they changed their morphology toward cluster 3. As the complex go/no-go odor learning task led to an increase in spine density, this might reflect the addition of new spines. If this is the case, it would indicate that a complex odor learning task leads to the formation and stabilization of new spines that predominantly belong to cluster 5. On the other hand, sensory deprivation decreased spine density with no noticeable changes in the spine clusters, indicating that the decrease was rather homogenous across the various clusters. At the first sight, the homogenous decrease in the density of spines across the various clusters seem at odds with our previous results showing that spines with spine head filopodia-like protrusions (SHF) are selectively preserved after sensory deprivation (Breton-Provencher et al., 2016). This can be explained by the fact that, first, SHF are very motile and appear and disappear very rapidly within a few minutes (Breton-Provencher et al., 2016). They can be easily observed in time-lapse imaging studies. However, due to their rapid dynamics, they are under-represented in the fixed tissue that we used for our current analysis. Second, SHF were also very thin and were more difficult to reconstruct with our pipeline. Future studies to perform 3D reconstruction and clustering analyses of spines derived from in vivo two-photon imaging studies in the OB, as we did previously (Breton-Provencher et al., 2016), combined with further improvements to the 3D reconstruction pipeline, will help address this issue.

In addition to learning-induced changes in the morphometric properties of spines and/or their density, our observations of marked changes in the morphology of the spines of the two control groups performing simple and complex tasks without associating the odor with a water reward (i.e., learning) were particularly intriguing. Almost all the clusters, except cluster 1, were affected. Clusters exhibiting larger changes included cluster 3, which decreased from 17.7% following the simple odor task to 2.17% following the complex go/no-go task, and cluster 4, which showed opposite changes (24.5% following the simple odor go/no-go task vs. 42.5% following the complex go/no-go task). The almost complete disappearance of cluster 3, which was central to the simple odor learning-induced changes, following the go/no-go complex task highlighted once again the different levels of structural modification of adult-born GCs in response to environmental challenges. This also indicates that the structural plasticity of adult-born GCs might be triggered not only in response to odor learning but also in response to the difficulty of the operating task, the duration of the odor stimulation, or the complexity of the odor mixtures. Future studies to test the role of passive exposure of mice to complex odor mixtures without exposing them to the go/no-go operational task will help resolve these issues.

### Limitations of the study

One of the limitations of our study was that we tailored our 3D reconstruction and computational pipeline to confocal images. We selected this approach given that confocal microscopy is widely used and enables the rapid testing and adaptation of our pipeline to user-specific needs. However, confocal microscopy does not provide sufficient resolution to resolve intricate details of spine morphology. Super-resolution microscopy, such as stimulated emission depletion (STED) microscopy, has recently shown that only a few, if any, stubby spines can be observed in organotypic slices of the hippocampus. The lower resolution of confocal and two-photon microscopy may lead to the classification of short-necked mushroom spines into the stubby spine category (Tønnesen et al., 2014). It is thus possible that spines in cluster 1 of the present study were short-necked mushroom spines. It should be mentioned, however, that independent of the type of spines in cluster 1, our clustering analysis based on various morphometric properties showed that spines in this cluster had distinctive characteristics compared to the other clusters.

Another limitation of our study was the lack of functional and ultrastructural signatures of clustered spines. Future studies using electron microscopic analyses of spines in a particular cluster and a functional assessment of their properties by combined glutamate uncaging and Ca2^+^ imaging will provide a better understanding of the functions of the spines in each specific cluster and how they are involved in the functioning of the bulbar network.

Lastly, it should be noted that GCs are a very heterogenous population of neurons that are characterized by the expression of distinct neurochemical markers and spatio-temporal assignments in the bulbar network (Batista-Brito et al., 2008, Malvaut et al., 2017, Hardy et al., 2018). Each GC subtype may be involved in a different odor behavior (Hardy et al., 2018). In the present study, we did not determine whether learning or sensory stimulation-associated changes in spine density and morphometric properties were linked to specific neurochemical signatures of adult-born GCs. Future studies that combine genetic targeting of different subtypes of adult-born GCs and computational assessments of learning-induced morphometric changes in spines will help address these issues.

Although all these limitations will pave the way path for future studies that will broaden our understanding of the structuro-functional properties of spines, the present study revealed that there is a vast panoply of distinct types of structural plasticity in adult-born GCs that are tailored in response to environmental stimuli and learning experiences.

## Methods

### Animals

Adult (>2-month-old) male C57BL/6 mice (Charles River) were used for all experiments. The experiments were performed in accordance with Canadian Guide for the Care and Use of Laboratory Animals guidelines and were approved by the Animal Protection Committee of Université Laval. The mice were kept in groups of 4-5 on a 12-h light/dark cycle in a temperature-controlled facility (22°C), with food and water ad libitum. The animals that underwent the go/no-go odor discrimination procedure were partially water-deprived, individually housed, and kept on a reverse light-dark cycle for 7-10 days before beginning the experiments. Animals were randomly assigned to the various experimental groups.

### Stereotaxic surgeries

We used a GFP-expressing lentiviral vector (LV-EF1α-puro, SignaGen Laboratories, SL100269) to label neuronal precursors and image GC dendritic spines. Mice were stereotaxically injected in the RMS under isoflurane anesthesia (2-2.5% isoflurane, 1 L/min of oxygen) and were kept on a heating pad during the entire surgical procedure. The following coordinates (with respect to the bregma) were used to target the RMS: anterior-posterior (AP) 2.55, medio-lateral (ML) 0.82, and dorso-ventral (DV) 3.15. The mice were allowed to recover on a heating blanket in a clean cage before returning to the housing room.

### Sensory deprivation

Two weeks after the viral injections, a group of animals was unilaterally sensory deprived. Occlusion tubes were made using polyethylene tubing (PE50, I.D. 0.58 mm, O.D. 0.965 mm; Becton Dickinson), with the center blocked using a tight-fitting Vicryl suture knot (3-0; Johnson & Johnson). The 5-mm-long petroleum jelly-coated plugs were inserted in the left nostril of the mice under isoflurane anesthesia. The mice were sacrificed by intracardiac perfusion 14 days later (i.e., 4 weeks post-viral injection).

### Go/no-go olfactory discrimination learning

Approximately 4-5 weeks after the injection of the GFP-expressing lentivirus, the mice were partially water-deprived to reach 80%-85% of their initial body weight prior to starting the go/no-go training. They first underwent training sessions to habituate themselves to the olfactometer chamber and learn how to get a water reward. The mice were trained using 20 trials, with no exposure to an odor, to insert their snouts into the odor sampling port and lick the water port to receive a 3-µL water reward. The two ports were located side-by-side. The reward-associated odor (S+) was then introduced. Inserting the snout into the odor sampling port broke a light beam and opened an odor valve. The duration of the opening was increased gradually from 0.1 to 1 s over several sessions, and mice with a minimum sampling time of 50 ms were given a water reward. The mice usually completed this training after one or two 30 min sessions. Once they had successfully completed this training step, they were subjected to the go/no-go odor discrimination test. Prior to being introduced to the non-reinforced odor (S-), the mice underwent an introductory S-session consisting of exposure to S+ for 30 trials. If the success rate was at least 80%, the discrimination task was begun. The mice were then exposed randomly to S+ or S-, and the percentage of correct responses was calculated for each block of 20 trials. If a mouse licked the water port after being exposed to S+ (hit) and did not lick the water port after being exposed to S- (correct rejection), this was recorded as a correct response. A false response was recorded if the mouse licked the water port after being exposed to S-or if it did not lick the water port after being exposed to S+. A mouse was considered successful if it reached a criterion score of 80%. The control group underwent the same go/no-go procedure but received the water reward independently of the odor used (S+ or S-). Mice from both groups underwent one of the two versions of the task.

The simple task involved 0.1% octanal (Sigma Aldrich) diluted in 99.9% mineral oil (Sigma Aldrich) as S+ and 0.1% decanal as S-. The complex version of the task involved two mixtures of similar odorants: 0.6% (+)-limonene (Sigma Aldrich) + 0.4% (−)-limonene (Sigma Aldrich) (S+) diluted in 99.9% mineral oil vs. 0.4% (+)-limonene, + 0.6% (−)-limonene (S−) diluted in 99.9% mineral oil. The mice from the two groups were sacrificed by intracardiac perfusion 1 h after completing the task.

### Immunohistochemistry

The mice were deeply anesthetized using an intraperitoneal injection of sodium pentobarbital (12 mg/mL; 0.1 mL per 10 g of body weight) and were intracardially perfused with 0.9% NaCl followed by 4% paraformaldehyde (PFA). The brains were collected and were post-fixed overnight in 4% PFA. Horizontal 100-μm-thick OB sections were cut using a vibratome (Leica) and were incubated overnight with a chicken anti-GFP primary antibody (Aves Labs, GFP-1020, 1:1000) diluted in 1% PBS supplemented with 0.5% Triton X-100 and 4% milk. For animals that underwent the sensory deprivation procedure, some slices were incubated overnight with a mouse anti-tyrosine hydroxylase (TH) primary antibody (Immunostar, ref. 22941, 1:1000) to compare deprived and control OBs. TH labelling is commonly used to assess sensory deprivation efficiency, with occlusion resulting in a 50% decrease in the TH signal in the ipsilateral hemisphere (Bastien-Dionne et al., 2010). The corresponding secondary antibodies were then used. Fluorescence images were acquired using an inverted Zeiss microscope (LSM 700, AxioObserver) with a 63X oil immersion objective (Plan Apochromat, NA: 1.4). To optimize image acquisition, we used a resolution of 1024 x 1024 pixels and an optical sectioning of 0.35 μm. Confocal microscopy was chosen because it is a widely used technique to visualize spines. For reproducibility purposes and for the robustness of the analysis, the same imaging parameters were used for the entire study.

### Dendritic and dendritic spine reconstructions

First, animals and experiments were anonymized to avoid any bias during extraction. Confocal images of GC spines collected from the different experimental groups were pre-processed with the DAMAS deconvolution algorithm (Brooks and Humphreys, 2006, Dougherty, 2005) using the Iterative Deconvolve 3D plugin in ImageJ 1.53c (Schneider et al., 2012). The images were then segmented using a Morphological Active Contours without Edges algorithm (Alvarez et al., 2010, Marquez-Neila et al., 2013, Chan and Vese, 2001) with the convergence set at 100 iterations and the parameter µ set at 1, which influences the smoothing and depends on the complexity of the segmented object. The other two parameters, λ_1_ and λ_2_, were set at 1 and 9, respectively, as these parameters are related to the ratio between intensity and background. These parameters were determined empirically to increase the signal-to-noise ratio. After segmentation, some images had visual artifacts in the form of single black pixels (value=0) and were removed with a script. In this script, for each black pixel, if the pixel was surrounded by at least 8 pixels with a value >0, the value of the black pixel was set as the mean of the surrounding pixels. The processed images were converted into meshes using the marching cubes algorithm (Lewiner et al., 2003, Lorensen and Cline, 1987). The spacing used in the algorithm was [0.24, 0.05, 0.05], which corresponds to the pixel size for axes z, x, and y. The z axis was corrected for spherical aberration (Kashiwagi et al., 2019). A polygonal mesh was generated using triangles to reproduce the surface of the spine. Each triangle was composed of vertices and was connected by edges. Before extracting the spines, a custom optimization process was performed using the following steps:

1. Creation of a box around the mesh
2. Determination of the target length based on the diagonal of the mesh box
3. Choice of the target level of detail (normal was used)
4. Removal of degenerated triangles (collinear triangles)
5. Removal of isolated vertices corresponding to vertices not connected to a face or edge
6. Removal of self-intersecting edges and faces
7. Removal of duplicated faces
8. Second removal of isolated vertices
9. Calculation of outer hull volume to check for any reconstruction problems with the mesh
10. Removal of obtuse triangles (angle >179°)
11. Removal of duplicated faces and vertices
12. Assessment of a potentially broken mesh: if a mesh was actually broken, the process was repeated starting at step 3 using a lower level of detail

After this reconstruction, the spines were extracted manually from the dendrite using Meshlab software (Cignoni et al., 2008).

### Measurement of morphometric features

To measure the morphometric features of the reconstructed spines, the hole left from the separation of the spine from the dendrite was used to calculate the position of the spine base center (S_bc_). Graph theory was used to assess the number of neighbors of each vertex. Because the vertices with the lowest number of neighbors were closest to the spine base with only three neighbors, the spine base center can be defined as the mean coordinate of these vertices.

#### Length (L)

The Length between each vertex (N) of the mesh in three dimensions (x, y, z) and the spine base center (S_bc_) is first calculated:

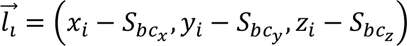

Then, a new mathematical subset is defined with the 5% longest distance (*n*). The final length is determined as the mean of these lengths:

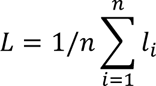

#### Surface (S)

The spine Surface area is the sum of all the triangle surfaces (s):

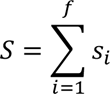

A triangle surface (s) is defined for every face (f) of the spine. For each triangle (*i*) with summits A, B, and C:

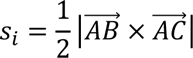

#### Volume (V)

The spine Volume is the sum of the entire tetrahedron volume (v):

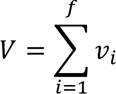

Each tetrahedron is defined by 4 vertices corresponding to triangles A, B, and C, and the spine center (S_C_) for all three-space coordinates (x, y, z). The tetrahedron volume is then calculated:

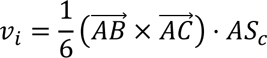

#### Hull Volume (HV)

The Hull Volume represents the smallest convex set containing all the vertices of a geometric shape. The calculation of the Hull Volume is performed using the built-in function of the Python package trimesh 3.7.10, which is based on the qhull algorithm (Barber et al., 1996).

#### Hull Ratio (HR)

The Hull Ratio is determined as:

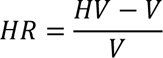

It can represent the complexity of the structure. The higher the score, the more complex the mesh.

#### Average Distance (AD)

The Average Distance represents the mean distance between each vertex (N) and the spine base center.

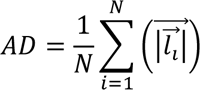

#### Coefficient of Variation in Distance (CVD)

The Coefficient of Variation in Distance represents the standard deviation of the variation of distance between the base of the spines and each vertex.

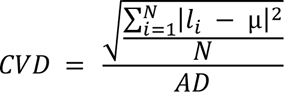

A high CVD is correlated with a large variation between the distance of the spine base center and the vertices. A high CVD often leads to mushroom-like spines while a small CVD leads to small stubby or small thin spines.

#### Open Angle (OA)

The Open Angle corresponds to the average angle between the spine axis and each vertex vector. The spine axis was defined as the vector crossing the gravity center (G_c_) and the mesh base center (S_bc_). The vertex vector is defined as the vector (l_i_) crossing the vertex and the mesh base center.

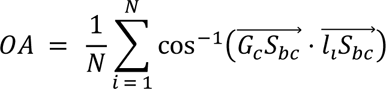

Large head spines are associated with a higher OA value, and thin spines with a low OA value.

#### Mean Curvature (MC)

The Mean Curvature corresponds to the mean of the curvature of each face of the mesh. It was calculated using trimesh (Cohen-Steiner and Morvan, 2003) and was then normalized to the surface.

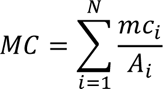

#### Mean Gaussian Curvature (GC)

The Mean Gaussian Curvature corresponds to the mean of the gaussian curvature of each face of the mesh. The main difference between GC and MC is that GC is quadratic while MC is linear. It was calculated using trimesh based on (Cohen-Steiner and Morvan, 2003) and was then normalized to the surface.

### Dimension reduction, clustering, and statistical analysis

The correlation matrix was calculated, and a Principal Component Analysis (PCA) was performed using the Python package scikit-learn 1.0.2. As the spine features dataset was not susceptible to the curse of dimensionality (Köppen, 2000, Verleysen and François, 2005), we used PCA to reduce the number of features from five to three independent components. Altogether, these three Principal Components (PC) contained 85.4% of the variance in the dataset. The clustering was performed using the K-Means algorithm (Arthur and Vassilvitskii, 2006) included in scikit-learn 1.0.2. The number of clusters was determined and was validated using three different scores: the Calinski-Harabz score (Caliński and Harabasz, 1974), the Silhouette score (Rousseeuw, 1987), and the Elbow score (Yuan and Yang, 2019). After clustering, the significance between the percentage of each cluster was determined using a two-independent proportion Agresti-Caffo test, which is available in statsmodels (Agresti and Caffo, 2012). *P* values <0.05 are considered statistically significant (\**p*<0.05, \*\**p*<0.01, \*\*\**p*<0.001). The Python package seaborn 0.12.1dev0 was used to generate the KDE maps. The Python package pingouin 0.5.3 was used to calculate Cohen’s term d. An effect lower than 0.01 is considered very small while an effect lower than 0.20 is considered small.

## Material Availability

Further information and requests for resources and reagents should be directed to and will be fulfilled by the co-corresponding authors, Dr. Armen Saghatelyan and Dr. Simon V. Hardy.

## Acknowledgments

This work was supported by a Canadian Institute of Health Research (CIHR) grant to A.S., National Science and Engineering Research Council of Canada (NSERC) grants to S.V.H. and A.S., and a Le Fonds de recherche du Québec – Nature et technologies (FRQNT) team grant to A.S. and S.V.H.

## Author contributions

A.F. performed the computational analyses of the spines of adult-born neurons. V.C. and S.M. performed the experiments and acquired the images. S.V.H. and A.S. supervised the computational and experimental work, respectively. All the authors discussed the data, and A.F., S.M., A.S., and S.V.H wrote the manuscript.

